# Asymmetric HIV-1 envelope trimers bound to one and two CD4 molecules are intermediates during membrane binding

**DOI:** 10.1101/2022.12.23.521843

**Authors:** Wenwei Li, Elizabeth Nand, Zhuan Qin, Michael W. Grunst, Jonathan R. Grover, Julian W. Bess, Jeffrey D. Lifson, Michael B. Zwick, Hemant D. Tagare, Pradeep D. Uchil, Walther Mothes

## Abstract

Human immunodeficiency virus 1 (HIV-1) infection is initiated by binding of the viral envelope glycoprotein (Env) to the cell-surface receptor CD4. Although high resolution structures of Env complexed with soluble domains of CD4 have been determined, the binding process is less understood on native membranes. Here, we apply cryo-electron tomography (cryo-ET) to monitor Env-CD4 interactions at membrane-membrane interfaces formed between HIV-1 and CD4-presenting virus-like particles. Env-CD4 complexes organized into clusters and rings, bringing opposing membranes closer together. Additionally, Env-CD4 clustering was dependent on capsid maturation. Subtomogram averaging and classification revealed that Env bound one, two, and finally three CD4 molecules, upon which Env adopted a partially open state. Our data indicate that asymmetric HIV-1 Env trimers bound to one and two CD4 molecules are detectable intermediates during virus binding to host cell membranes, which likely has consequences for antibody-mediated immune responses and vaccine immunogen design.

## Introduction

The human immunodeficiency virus 1 (HIV-1) begins infection of CD4^+^ T cells when the trimeric HIV-1 envelope glycoprotein (Env) binds to the cell-surface receptor CD4^1-4^. CD4 binding induces conformational changes within the gp120 subunit of Env that allow subsequent engagement of co-receptors CCR5 or CXCR4. Receptor and co-receptor engagement triggers conformational changes in the Env gp41 subunits which drive fusion of the virus and host cell membranes. The first structural insights into HIV-1 Env interactions with the CD4 receptor were made possible by the development of a gp120 core lacking variable loops that was crystalized with the D1-D2 domains of CD4 and the Fab of the co-receptor mimicking antibody 17b^5^. The next major breakthrough in understanding Env structure came with the development of stable, soluble Env trimers (trimers containing an S-S disulfide bridge and an I559P mutation, or SOSIPs) suitable for structural studies^6-9^. Studies of SOSIP trimers revealed that in the closed state, the variable loops (V) 1 and 2 form the apex of the Env trimer. CD4 binding results in an ∼40Å displacement of the V1V2 loop to expose the underlying co-receptor binding site in the V3 loop; the displaced V1V2 loop aligns with the CD4 D1D2 domains^10-13^. The bridging sheet undergoes extensive rearrangements upon CD4 binding, with the inner helix of the heptad repeat 1-N (HR1N) domain extending toward the opposing membrane^11,13^. The insights gained from soluble trimers have been confirmed with full-length Env proteins solubilized from membranes using detergent^14-16^. Further information about the co-receptor binding step of Env were made by embedding coreceptor CCR5 in lipid nanodiscs and determining the structure of the complex of soluble gp120, soluble CD4 (D1-D4), and CCR5^17^. In this structure, the V3 loop of gp120 reached into a cavity formed by the CCR5 helices, and the N-terminus of CCR5 was bound to the gp120 bridging sheet exposed by CD4 binding. As such, binding of CCR5 did not induce additional allosteric changes in gp120, but rather brought the CD4-bound gp120 closer to the lipid bilayer mimicked by the nanodisc. Although completely open and partially open CD4-bound Env conformations have been observed with SOSIP trimers^10,11^, whether the partially open Env trimer conformations are true intermediates is unknown.

These structural insights into HIV-1 entry were achieved using stabilized soluble trimer proteins and the soluble D1D2 domains of CD4. Further advances have been made using HIV-1 Env trimers in native membranes interacting with soluble receptors. Directly imaging virus particles by cryo-electron tomography (cryo-ET) and subsequent subtomogram averaging has allowed a characterization of the native Env trimer and the Env trimer opened by soluble CD4 initially at ∼20Å^18^ and more recently at ∼9-10Å^19,20^. These structures largely confirm the high resolution Env structures obtained with soluble trimers and soluble ligands, including the extension of the inner helix of the HR1N upon binding of sCD4^19^. Furthermore, insights into the behavior of HIV-1 Env molecules on the surface of virus particles have been gained from single molecule FRET studies in which a donor and acceptor fluorophore were introduced into a single gp120 subunit of a trimer^21^. These studies indicated that individual protomers are dynamic and have spontaneous access to open conformational states including the CD4-bound state^21,22^.

Engineering of trimers that can only bind one or two CD4 molecules suggested that a necessary intermediate FRET state in the opening of Env corresponds to an asymmetric trimer in which only one CD4 molecule binds the trimer^22^. Additionally, a structure of a soluble trimer mutationally prevented from opening has been observed to bind only a single CD4^23^. Finally, the interaction of HIV-1 Env molecules with CD4 and coreceptor molecules have been studied in living cells by combining super-resolution localization microscopy with fluorescence fluctuation spectroscopy imaging^24^. These studies suggested that HIV-1 entry is initiated by Env binding to a single CD4, followed by recruitment of additional CD4 molecules and a dimer of coreceptor molecules. Structures of native HIV-1 Env trimers bound to one or two CD4 molecules have not been observed in membranes.

Studying how HIV-1 Env in native membranes interacts with soluble receptors has advanced our understanding of HIV-1 entry. However, it remains unknown how membrane embedded full-length CD4 receptor engages the Env trimer. Thus, a complete understanding of the HIV-1 entry process requires studies of Env-receptor interactions on native membranes. Here we directly visualize the interactions between HIV-1 Env in virions and native membrane-bound CD4 by cryo-ET. We developed an experimental system that generates membrane-membrane interfaces presenting Env and CD4 by mixing HIV-1_BaL_ particles with receptor-carrying virus-like particles (VLPs). This approach generates a high frequency of Env-CD4 complexes at membrane-membrane interfaces, allowing for quantitative analysis by cryo-ET and subtomogram averaging. Using this mixed virus particle system, we observed that Env trimers cluster and organize into rings, and the patterns of clustering correlate with decreasing distances between membranes at the interfaces. We also observed that Env clustering depends on capsid maturation, as demonstrated by both cryo-ET and a fusion assay based on split nanoluciferase (nanoluc) complementation. Following subtomogram averaging of CD4-bound Env trimers, we solved an ∼16Å structure of Env bound to three native, membrane-bound CD4 receptors. Classification of the distance between opposing membranes revealed that when membranes are farther apart, an Env trimer engaged a single CD4 molecule. As the opposing membranes approached each other, Env trimers bound two or three CD4 molecules. The V1V2 loop projected outward in the CD4-bound protomers while the unbound protomers showed heterogenous conformational states with unclear V1V2 loop density. These data indicate that asymmetric HIV-1 Env trimers with a single and two CD4 molecules bound are detectable intermediates during virus binding to membranes.

## Results

### Env-CD4 interactions captured in an experimental system compatible with cryo-ET

We used cryo-ET to characterize membrane embedded HIV-1 Env-CD4 interactions *in situ*. Cryo-ET imaging is constrained by the depth of the vitreous ice layer, ∼500 nm. Therefore, we developed an experimental system that is uniform and presents observable Env-CD4 complexes at high frequency by using viral particles, which are the ideal size for cryo-ET imaging (∼150 nm). In this system, we utilized HIV-1_BaL_ particles produced from chronically infected SupT1 T cells, which have a high density of Env trimers making them suitable for use in cryo-ET studies^18,19^. Since they are highly infectious, they were inactivated with AT-2 before imaging (see methods); AT-2 inactivation does not interfere with CD4 binding or fusion activity of these particles^20,25^. Using a Murine Leukemia Virus (MLV) GagPol we also produced virus-like particles that carry the CD4 receptor, with or without co-receptor CCR5 (MLV-CD4 particles). HIV-1_BaL_ particles and MLV-CD4 particles were mixed and plunge-frozen for imaging by cryo-ET (Fig. 1a). HIV-1 and MLV have distinct capsid morphologies, meaning that the two types of particles could be visually distinguished in the cryo tomograms (Fig. 1b-c). The different capsid morphologies also allowed for the identification of membrane-membrane interfaces that were the consequence of Env-CD4 interactions vs. those that occurred due to spatial proximity (Fig. 1d). While individual Env molecules could be visualized on HIV-1_BaL_ particles, the smaller CD4 molecules were not easily identifiable on MLV particles. However, upon binding to Env at membrane-membrane interfaces, the CD4 receptor molecules could be visualized (Fig. 1e-f).

**Fig. 1.**
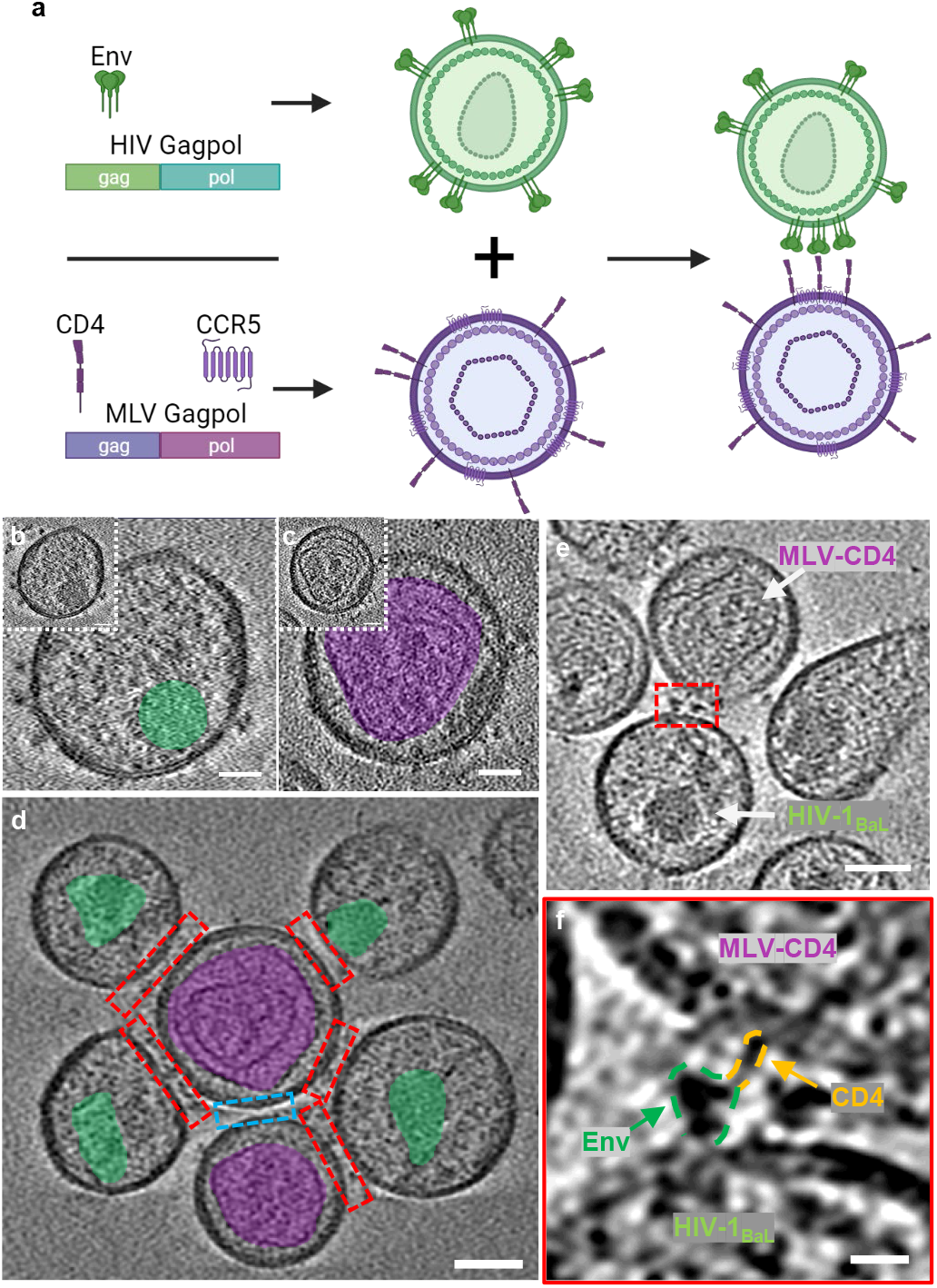
Env-receptor interactions captured in an experimental system compatible with cryo-ET. **a** Schematic illustration of the mixed viral particle system with HIV-1_BaL_ virions presenting Env and MLV VLPs presenting CD4 with or without co-receptor. **b** Representative image of HIV-1_BaL_ virus with Env on its surface. The capsid is highlighted in green. HIV-1_BaL_ was inactivated with AT-2, resulting in less electron-dense capsids. Scale bar = 25 nm. **c** Representative images of MLV VLP carrying CD4 on its surface. The MLV capsid is highlighted in purple. Scale bar = 25 nm. **d** A representative image of membrane-membrane interfaces (red boxes) in cryo-tomograms. Different capsid structures allow for identification of membrane-membrane interfaces as opposed to interfaces that do not have Env-CD4 interactions (blue box). Scale bar = 50 nm. **e** A representative tomogram with the membrane-membrane interface highlighted in red. Scale bar = 50 nm. **f** Enlarged image of the membrane-membrane interface in panel e. Env-CD4 interactions are visible in raw tomograms. Scale bar = 10 nm.

### HIV-1 Env binding to CD4 induces Env clustering and ring formation at membrane-membrane interfaces

Patterns of Env clustering became apparent in the cryo-ET tomograms, with small clusters, large clusters, and rings of Env trimers being observed at membrane-membrane interfaces (Fig. 2a-d). Quantification of these clusters revealed that the number of Env trimers increased from small clusters to large clusters and rings (Fig. 2e). Additionally, the distance between the opposing membranes changed with small clusters having the greatest membrane distance, large clusters having an intermediate distance, and rings having the smallest distance (Fig. 2f). To test if the presence of CD4 correlated with clustering into the observed patterns, all arc distances between two Env trimers on the surface of virus particles were calculated in the presence and absence of MLV-CD4 particles. The presence of MLV-CD4 VLPs increased the frequency of short arc distances between Env trimers on the surface of HIV-1 virions (Fig. 2g). This indicates that a subpopulation of trimers are drawn into clusters at membrane-membrane interfaces in a CD4-dependent manner.

**Fig. 2.**
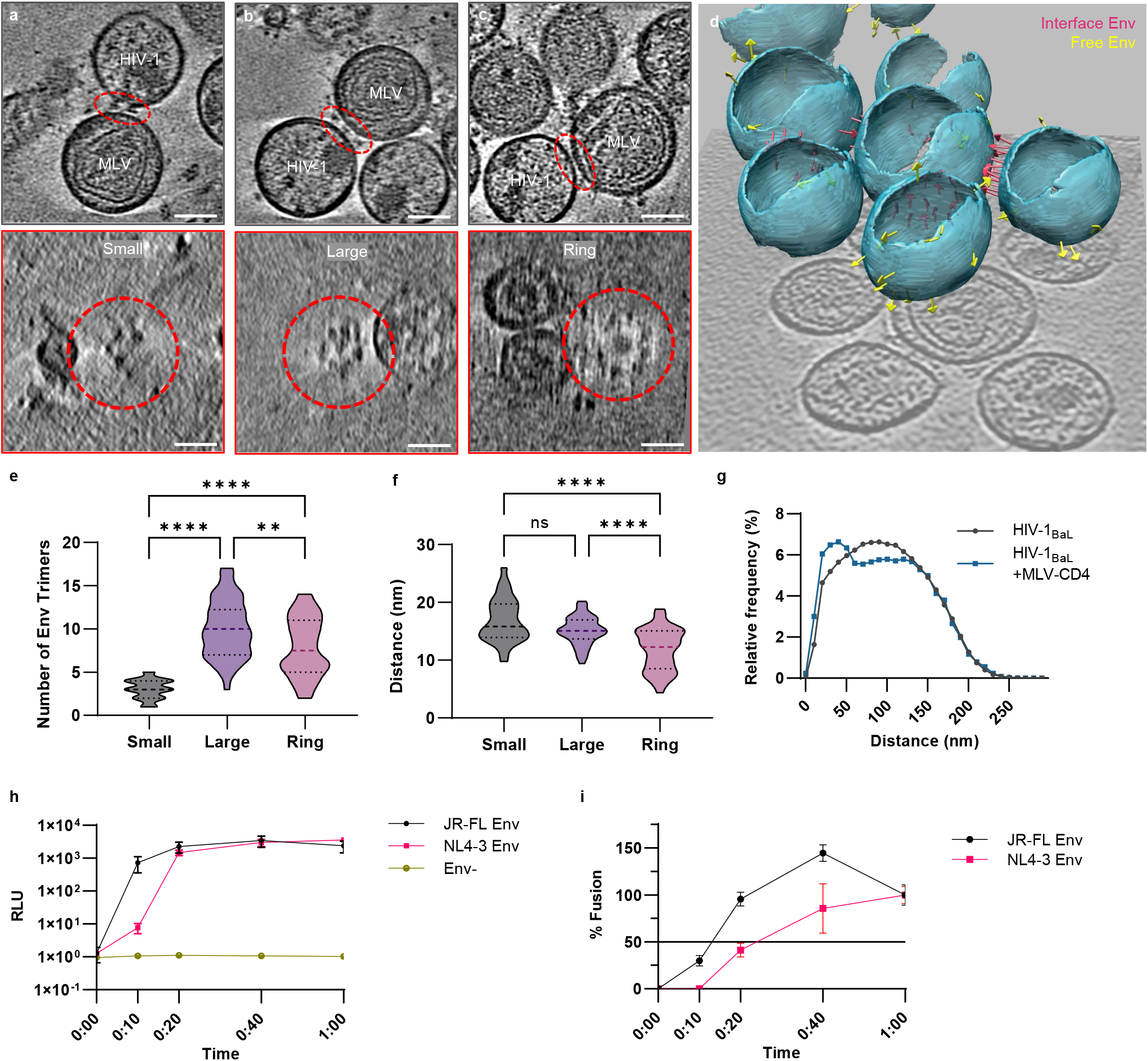
HIV-1 Env binding to CD4 induces Env clustering and ring formation at membrane-membrane interfaces. **a-c** Representative tomograms of small, large, and ring cluster formations. Interfaces are highlighted in red. The lower panel shows the top-down view of each interface, revealing the Env clustering. Scale bars = 50 nm. **d** Three-dimensional representation of viral particles in a representative tomogram. Pink arrows represent Env trimers at a membrane-membrane interface. Yellow arrows represent free Env trimers on the surface of HIV-1_BaL_. **e** Violin plot of the number of Env trimers present at interfaces in each clustering pattern. Small cluster mean = 3.0 ± 1.1. Large cluster mean = 10.0 ± 3.4. Ring cluster mean = 7.9 ± 3.4. P-values ** = 0.0024, **** = <0.0001. **f** Violin plot of the distance between membranes at interfaces in each clustering pattern. Small cluster mean = 16.9 nm ± 3.8 nm. Large cluster mean = 15.1 nm ± 2.4 nm. Ring cluster mean = 11.9 nm ± 3.6 nm. P-values **** = <0.0001. **g** Histogram of the arc distances between Env trimers on the surface of HIV-1_BaL_ particles alone (black) and HIV-1_BaL_ particles mixed with MLV-CD4 VLPs (blue). **h** Fusion kinetics of HIV-1 VLPs carrying Env_JR-FL_ (black) and Env_NL4-3_ (red). Two independent experiments were performed in triplicate for each Env strain and data sets were combined and plotted. Error bars are SD. **i** Data in panel (h) shown as percent of total fusion at 1 hr. All 1 hr time point data for each Env strain was averaged and set to 100%. 50% fusion is marked on the graph. Error bars are SD.

Env clustering at membrane-membrane interfaces has implications for Env stoichiometry during fusion, with strains that require more Env trimers for fusion (such as NL4-3) expected to be more dependent on Env clustering than those that require fewer Env trimers for fusion (such as JR-FL) ^26^. This dependence on Env clustering can be tested using fusion kinetics. As such, we developed a real time virus-cell fusion assay using split nanoluc complementation to test fusion kinetics of HIV-1_NL4-3_ and HIV-1_JR-FL_ strains. The results showed an ∼10-minute delay in fusion for the Env_NL4-3_ relative to the Env_JR-FL_ (Fig. 2h-i). The lag time in fusion kinetics observed with Env_NL4-3_ is consistent with additional time needed for 5 Env trimers, the approximate number of trimers necessary for HIV-1_NL4-3_ fusion, to form a cluster. The faster fusion kinetics of Env_JR-FL_ is also consistent with these data, as HIV-1_JR-FL_ is hypothesized to require only 1-2 trimers for fusion^26^.

### HIV-1 Env clustering and fusion kinetics are dependent on capsid maturation

HIV-1 Env mobility on the surface of the virus particle has been observed to be dependent on capsid maturation and the cytoplasmic tail (C-tail) of Env^27-29^. In the immature capsid, Env is laterally immobilized by the C-tail tethering to the underlying immature matrix (MA). During maturation, MA is cleaved from the capsid, allowing Env to move laterally on the surface of the viral particle^28,29^. To test if Env clustering induced by CD4 binding is impaired in immature capsids, MLV-CD4 particles were incubated with HIV-1_BaL_ particles produced in the presence and absence of protease inhibitors Indinavir (IDV) and Ritonavir (RTV), then processed for imaging by cryo-ET. Immature HIV-1 capsids remain morphologically distinct from MLV capsids, allowing for reliable identification of HIV-1_BaL_ particles relative to MLV-CD4 particles (Fig. 3a). Quantification of tomograms revealed that membrane-membrane interfaces with immature capsids had fewer Env trimers (4.6 ± 2.5) than interfaces with mature capsids (7.5 ± 4) (Fig. 3b-c), although the membrane distance at interfaces showed no significant change between mature and immature capsids (Fig. 3d). Furthermore, the addition of MLV-CD4 particles to immature HIV-1_BaL_ particles did not decrease arc distances between two Env trimers as it did with mature HIV-1_BaL_ capsid particles, indicating that the immature capsid prevents Env clustering (Fig. 3e).

**Fig. 3.**
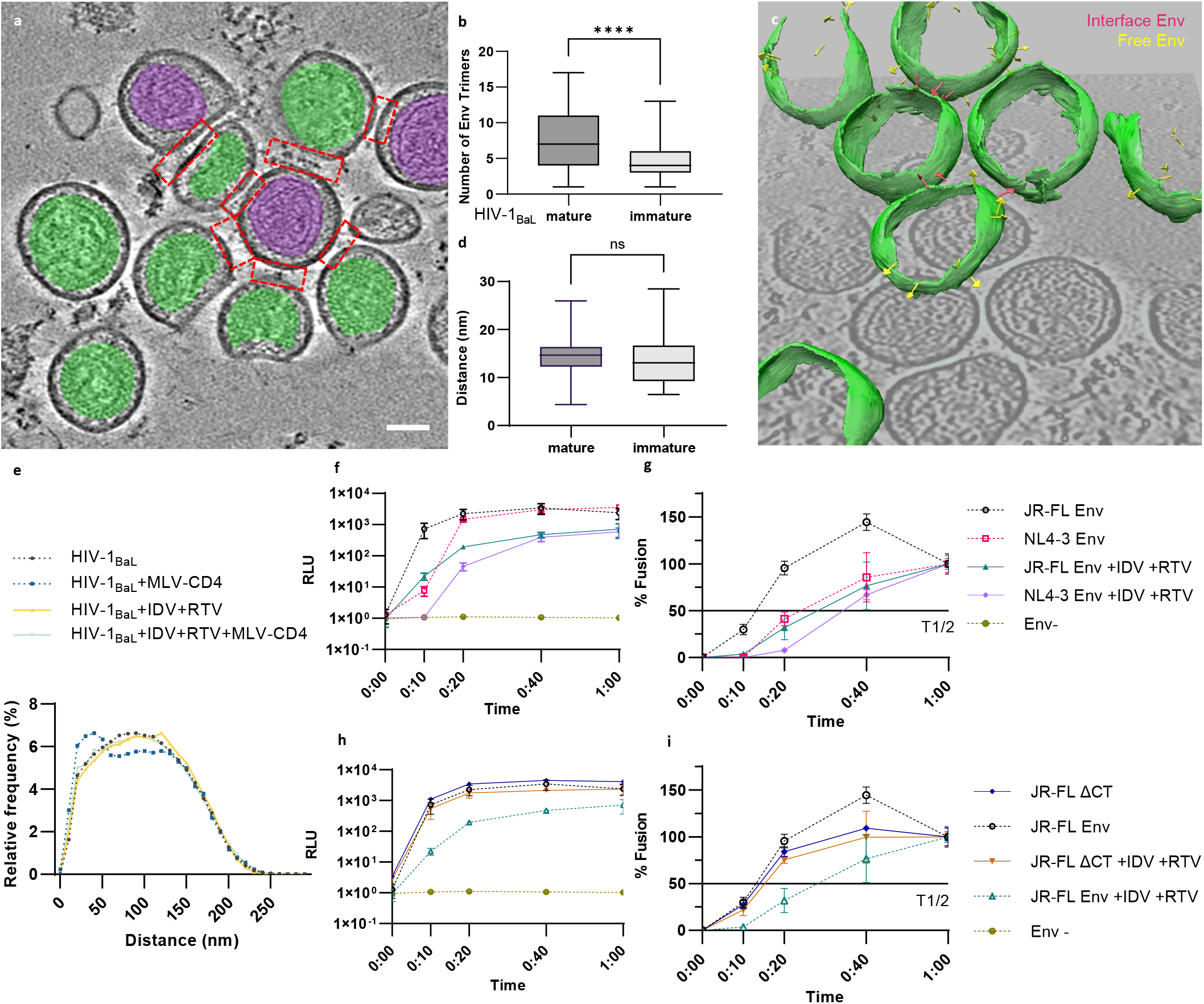
HIV-1 Env clustering and fusion kinetics are dependent on capsid maturation. **a** A representative tomogram showing immature HIV-1_BaL_ particles and MLV-CD4 VLPs. HIV capsids are noted in green and MLV capsids are noted in purple. Scale bar = 50 nm. **b** Box and whisker plot of the number of Env trimers present at membrane-membrane interfaces for mature and immature HIV-1_BaL_ particles mixed with MLV-CD4 VLPs. Mature HIV-1_BaL_ mean = 7.5 ± 4. Immature HIV-1_BaL_ mean = 4.6 ± 2.5. P-value **** = <0.0001. **c** Three-dimensional representation of immature HIV-1_BaL_ particles and MLV-CD4 VLPs in a representative tomogram. Pink arrows represent Env trimers at membrane-membrane interfaces. Yellow arrows represent free Env trimers on the surface of the HIV-1_BaL_ particles. **d** Box and whisker plot of the distance between membranes at membrane-membrane interfaces with mature and immature HIV-1_BaL_ particles mixed with MLV-CD4 VLPs. Mature HIV-1_BaL_ mean = 14.3 nm ± 3.8 nm. Immature HIV-1_BaL_ mean = 13.5 nm ± 4.6 nm. ns = not significant. **e** Histogram of the arc distances between Env trimers on the surface of mature (black) and immature (yellow) HIV-1_BaL_ particles alone, and mature (blue) and immature (green) HIV-1_BaL_ particles mixed with MLV-CD4 VLPs. **f** Fusion kinetics of immature HIV-1 VLPs carrying Env_JR-FL_ (green) or Env_NL4-3_ (lilac) compared to mature fusion kinetics in Fig. 2 (black and red dotted lines). Two independent experiments were performed in triplicate for each Env strain and data sets were combined and plotted. Error bars are SD. **g** Data in panel (f) shown as percent of total fusion at 1 hr. All 1 hr time point data for each Env strain was averaged and set to 100%. 50% fusion is marked on the graph. Error bars are SD. **h** Fusion kinetics of mature (blue) and immature (orange) HIV-1 VLPs carrying Env_JR-FL._Δ_CT_ compared to data from panel (f) of mature (black) and immature (green) HIV-1 VLPs carrying Env_JR-FL_ (dotted lines). Error bars are one SD. **i** Data in panel (h) shown as percent of total fusion at 1 hr. All 1 hr time point data for each Env strain was averaged and set to 100%. 50% fusion is marked on the graph. Error bars are SD.

To further confirm if the deficiency in Env clustering on immature capsids observed by cryo-ET was functionally relevant, fusion kinetics were measured for immature HIV-1_JR-FL_ and HIV-1_NL4-3_ VLPs. Immature VLPs were produced with Env_JR-FL_ or Env_NL4-3_ in the presence of IDV and RTV. With immature capsids, both strains showed delayed fusion kinetics compared to their untreated VLP counterparts with mature capsids (Fig. 3f, g). The delayed kinetics suggests that it takes additional time to accumulate sufficient numbers of Env trimers at membrane-membrane interfaces to facilitate fusion, thereby supporting that Env clustering plays a role in fusion. These experiments indicate that Env clustering likely contributes to productive infection not just for HIV-1 strains hypothesized to require more trimers for fusion (NL4-3), but also for strains that may require fewer trimers for fusion (JR-FL) ^26^.

To test if this phenotype is directly related to Env mobility rather than capsid maturation, fusion kinetics assays were performed with the C-tail deletion mutant (ΔCT) of HIV-1_JR-FL_ Env packaged on VLPs with both mature and immature capsids. The C-tail deletion of HIV-1_JR-FL_ Env rescued the delayed fusion kinetics imposed by immature capsid (Fig. 3h, i). This data suggests that Env clustering is precluded by interactions between matrix and the cytoplasmic tail of Env.

To further test the role of Env clustering during fusion, we used VLPs with high numbers of Env trimers. These VLPs were produced from a cell line that endogenously expresses HIV-1_ADA.CM_ Env that has lost a portion of its cytoplasmic tail (Env_ADA.CM.755*_)(truncated at residue 755) ^30^. We produced VLPs carrying Env_ADA.CM.755*_with either a wild type HIV-1 GagPol gene or HIV-1 GagPol with a protease knockout mutation (PR-) that precluded cleavage of the Gag precursor to liberate the Matrix protein. We imaged these VLPs on their own and in the presence of MLV-CD4 VLPs to test if Env clustering was still dependent on capsid maturation even with a very high number of Env trimers on the surface of the virion. Not surprisingly given the very high trimer density, a high number of Env trimers assembled at membrane-membrane interfaces. Furthermore and in agreement with our previous data, since the Env_ADA.CM.755*_ has a truncated cytoplasmic tail, the clustering was independent of PR-dependent capsid maturation (Extended Data Fig. 1a-d). Fusion kinetics of HIV-1 VLPs carrying Env_ADA.CM.755*_ were also independent of capsid maturation. This was the case for VLPs produced with the HIV-1 GagPol PR-mutant and those produced with WT GagPol in the presence of HIV-1 protease inhibitors IDV and RTV (Extended Data Fig. 1e). HIV-1_ADA.CM.755*_Env particles also provided an additional source of trimers to study the structure of Env-CD4 complexes at membrane-membrane interfaces.

### Subtomogram averaging and classification reveal intermediates with HIV-1 Env trimers bound to one, two, and three CD4 receptor molecules

Approximately 5700 subtomograms containing Env-CD4 complexes were manually picked from a total of 168 tomograms. Subtomogram averaging of these particles generated an ∼15 Å resolution density map of the HIV-1 Env trimer bound to three native, membrane-bound CD4 receptor molecules (Fig. 4a-d, Extended Data Fig. 2a). The trimers were clearly open with the gp120 density moving away from the central axis. The density for V1V2 loops projected outward in all three bound protomers of the Env trimer, which is consistent with previous high-resolution structures of soluble Env trimer bound to CD4^10,11^(Fig. 4b, d).

**Fig. 4.**
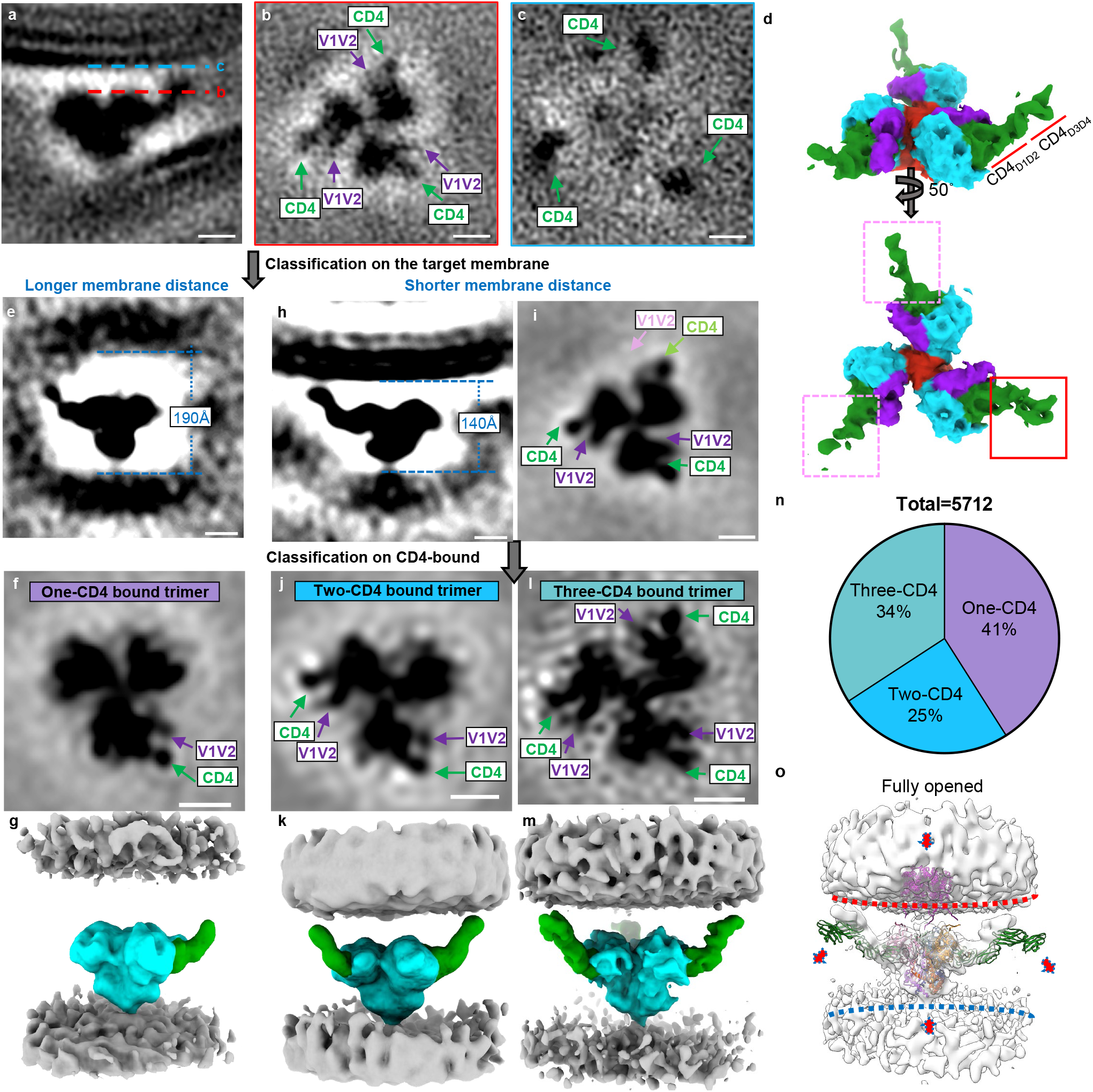
Subtomogram averaging and classification reveal intermediates with HIV-1 Env trimers bound to one, two, and three CD4 receptor molecules. **a-c** Side view (a) of subtomogram average of the Env-CD4 complex at membrane-membrane interfaces. Dotted lines indicate the positions of top-view sections (b) and (c). CD4 (b, c), and V1V2 loops (b) are indicated in green and purple, respectively. **d** Segmentation of Env-CD4 average structure. Side view (top) and top view (bottom) of Env (cyan) bound by three CD4 molecules (green). D1D2 and D3D4 domains of CD4 are labeled with red lines. V1V2 loops are shown in purple and gp41 is shown in red. The red box in the bottom panel shows the CD4 molecule with strong density in the average structure, while the pink dotted boxes show the two CD4 molecules with weaker densities. **e-g** Subclass average using longer membrane distance after focused classification on the target membrane. Side view (e), top view (f) and segmentation (g) of the structure are shown. The membrane distance (e) is indicated in blue. In the segmentation (g), Env is labeled in cyan and CD4 is labeled in green. Membranes are in grey. **h, i** Subclass average with shorter membrane distance after focused classification on the target membrane. Side view (h) and top view (i) are shown. The absence of the V1V2 loop density is labeled in pink and the distinctive CD4 is labeled in light green. **j-m** Further focused classification was performed from subtomograms in subclass average with shorter membrane distance (h, i). Subclass averages of Env bound to two CD4 (j, k) and three CD4 molecules (l, m) are shown with top views (j, l) and segmentations (k, m). Segmentations are colored as in panel (g). **n** Pie chart of proportion of different Env trimers bound to one, two, and three CD4 molecules. **o** Hypothetical conformational changes of the Env-CD4 complex required to engage the co-receptor. To overcome the steric constraints in CD4 molecules, gp120 shedding or membrane bending could facilitate the movement of Env towards the coreceptors embedded in target membranes. Scale bars = 5 nm in density maps.

We did not impose symmetry on the subtomogram averaging structure of the Env trimer bound to three CD4 molecules. One bound CD4 molecule featured stronger density than the other two in this structure and these two had the lowest resolution as shown by local resolution analysis, indicating the presence of structural heterogeneity in CD4 binding (Fig. 4d, Extended Data Fig. 2a). This, along with above observations that the membrane distance varied among different patterns of Env clustering, suggested that the structural heterogeneity was linked to the distance between membranes at membrane-membrane interfaces. We therefore performed subtomogram classification based on this membrane distance. The analysis revealed two subclasses; one with membranes farther apart (∼190 Å) where Env only engaged a single CD4 molecule (Fig. 4e, f, Extended Data Fig. 2b), and one with membranes closer together (∼140 Å) where Env engaged three CD4 molecules (Fig. 4h, i). Because heterogeneity remained in the densities of V1V2 loops and CD4 in the latter structure, we performed further subtomogram classification based on CD4 binding. This revealed an additional subclass of Env trimers with two bound CD4 molecules (Fig. 4j-m, Extended Data Fig. 2c). A total of 41% of Env trimers were bound to one CD4, 25% to two CD4, and 34% to three CD4 molecules (Fig. 4n). Thus, structures of Env bound to one, two, and three CD4 molecules are observable intermediates at membrane-membrane interfaces (Fig. 4g, k, m, Extended Data Fig. 2).

### Conformational changes in CD4 allow Env to approach the target membrane

We were able to resolve the structure of full-length, membrane-bound CD4 in the structure of Env bound to three CD4 molecules with a shorter membrane distance of ∼140 Å (Fig. 4m). In addition to the familiar structure of the D1D2 domains bound to Env, the D3D4 domains became clearly visible. The D3D4 domains of all three bound CD4 molecules were aligned with the target membrane. The observed angle between the D1D2 domains and the D3D4 domains is consistent with the previously observed flexibility in the D2-D3 hinge^31^. The alignment of the D3D4 domains and the flexibility of the D2-D3 hinge allowed CD4-bound Env to approach the target membrane. In the Env structure bound to one CD4 molecule, only the D1D2 domains of CD4 were resolved (Fig. 4g). To account for the observed membrane distance of ∼190 Å, CD4 must be in a fully extended conformation to be able to reach Env on the opposing membrane. Our data thus support a model where conformational changes in CD4 facilitate HIV-1 binding to membranes.

### HIV-1 Env trimers bound to one and two CD4 molecules are asymmetric with protomers adopting distinct conformational states

All structures of Env-CD4 complexes had well-defined densities of the bound CD4, with each Env protomer engaging CD4 displaying adjacent density for the outwardly projecting V1V2 loops (Fig. 4f, j, l). All Env-CD4 complexes, regardless of CD4 binding, were missing EM densities at the apex region, suggesting an open conformation where gp120 density moves away from the central axis (Extended Data Fig. 3a-d). This structure is similar to open conformations previously revealed by SOSIP trimers bound to soluble domains of CD4^10,11^. Interestingly, the free protomer in the trimer bound to two CD4 molecules lacked the density for the outwardly projected V1V2 loops. Although the resolution of our cryo-ET maps is not high enough to determine with certainty where the V1V2 loops sit, the conformational state is clearly distinct from the CD4-bound conformation (Fig. 4j, Extended Data Fig. 3f). Similar distinct conformational states are seen in the unliganded protomers of the Env trimer bound to one CD4 molecule, although some weak density for outwardly projected V1V2 loops remains visible (Extended Data Fig. 3e). This is likely attributable to remaining heterogeneity in the sample. Regardless, this analysis indicates that the HIV-1 Env trimers bound to one and two CD4 molecules are asymmetric open trimers with the CD4-bound protomer adopting an outward rearrangement of the V1V2 loops while the unliganded protomers remain in a distinct conformational state (Fig. 4f, j Extended Data Fig. 3a, b, e, f).

### Molecular dynamics flexible fitting suggests that the HIV-1 Env trimer bound to three CD4 molecules is in a partially open conformation

We wanted to test if existing atomic models of soluble Env trimers bound to soluble CD4 (D1D2) fit into our cryo-ET density of the Env trimer bound to three CD4 molecules^10,11^. We first generated a combined atomic model by superimposing the model of the gp120 monomer bound to a full-length CD4 molecule (D1-D4) and coreceptor CCR5 (PDB 6MET^17^) on the model of the soluble Env trimer bound to CD4 D1D2 domains (PDB 5VN3^10^). We docked this combined model into our density map using rigid body fitting. The combined model did not agree well with the cryo-ET structure (Extended Data Fig. 4a), so we performed Molecular Dynamics Flexible Fitting (MDFF) permitting full atom flexibility of the combined model within the cryo-ET density map (Extended Data Fig. 4b). There were large movements of the atomic model to fit the cryo-ET densities, particularly within the domains of CD4 where the center of mass shifted toward the target membrane as much as ∼19Å (Extended Data Fig. 4c, Extended Data Table 1, Extended Video 1). The domains of gp120 were also rearranged, moving closer to the central axis of Env trimer and producing a more closed/partially open structure (Extended Data Fig. 4d, Extended Data Table 1, Extended Video 1). Since the fully open state of PDB 5VN3 did not fit well into our cryo-ET density map, we next tested if a combined atomic model of a partially open soluble trimer bound with full-length CD4 receptors could fit in the cryo-ET densities (PDB 6CM3^11^). PDB 6MET was superimposed on PDB 6CM3 to create a second combined model, and this model fit well into the cryo-ET density (Extended Data Fig. 4e). MDFF resulted in an even better fit, though the gp120 still rearranged into a slightly more closed structure (Extended Data Fig. 4f-h, Extended Data Table 1, Extended Video 2). This analysis indicates that partially open trimers are observable intermediates during HIV-1 Env binding to membrane-bound CD4.

## Discussion

We applied cryo-ET to visualize the initial steps of HIV-1 entry where Env trimers engage CD4 receptor molecules residing in target membranes. Mixing HIV-1_BaL_ virions with VLPs presenting CD4 receptors provided an experimental system which generated a high frequency of Env-CD4 complexes, allowing us to quantitatively study their interactions in native membranes by cryo-ET. We observed HIV-1 Env trimers forming clusters and rings at membrane-membrane interfaces, and Env trimers engaging one, two, and three CD4 receptor molecules. As the distance between membranes at membrane-membrane interfaces decreased, structures of Env bound to increasing numbers of CD4 molecules became visible. This suggests that these images represent snapshots of a dynamic stepwise binding process.

The progression from clusters to rings at membrane-membrane interfaces is remarkably similar to the organization of SNAREs at membrane docking sites of synaptic vesicles^32^. It is possible that the patterns of Env-CD4 complexes observed at membrane-membrane interfaces may be a consequence of simple adhesion. However, both SNAREs and HIV-1 Env trimers are fusion machines that ultimately bring membranes together for fusion, suggesting that these observations are relevant for fusion.

The physiological relevance of Env trimer clustering at fusion sites is related to the minimum number of trimers required for fusion. Among enveloped viruses HIV-1 Env is unique; for some HIV-1 isolates, such as JR-FL, just one to two Env trimers may be sufficient to mediate fusion^26,33,34^, while the lab-adapted NL4-3 strain may require approximately five trimers for fusion^26^. While these studies applied computational models to arrive at these Env trimer numbers, live cell imaging of HIV-1 Env, CD4 receptor, and coreceptor molecules similarly indicated that as few as one to two Env trimers can recruit approximately four CD4 molecules and approximately two coreceptor molecules into a fusion-competent cluster^24^. The number of Env trimers needed for fusion is related to the fusogenicity of each Env strain; Env_NL4-3_ needs more trimers to reach the same fusogenicity that Env_JR-FL_ reaches with only one to two trimers^26^. Our fusion kinetics experiments are consistent with the additional time required for more Env trimers with lower fusogenicity, such as HIV-1_NL4-3_, to cluster at membrane-membrane interfaces. Furthermore, our experiments with immature capsids suggest that Env clustering is likely required even for isolates with high Env fusogenicity, such as HIV-1_JR-FL_.

The mixed viral particle system allowed us to visualize membrane-membrane interfaces containing Env-CD4 complexes at a high frequency. Acquiring 168 tomograms allowed us to identify 857 membrane-membrane interfaces containing ∼5700 Env-CD4 complexes for subtomogram averaging and subsequent classification. Our comprehensive analysis revealed that HIV-1 Env trimers engage a single CD4 when the two membranes are farther apart and bind a second and third CD4 as Env moves closer to the membrane. The conformational states of the Env trimers bound to one and two CD4 molecules are asymmetric, not just with respect to the inherent asymmetry in CD4 binding, but also with respect to the conformational state of each protomer within the trimer. The resolutions of our density maps were high enough to identify that the V1V2 loops project outward in all CD4-bound protomers. The density was less defined for the V1V2 loops in the unbound protomers of the Env trimers with one or two bound CD4 molecules. We expect that the V1V2 loops are still localized to the trimer apex in these protomers.

Previous smFRET studies performed by our group predicted the existence of asymmetry in the opening of the trimer^22^. Trimers engineered to bind only one CD4 molecule featured the neighboring unliganded protomers in an intermediate conformational state that was distinct from the unbound and CD4-bound conformations^22^. Importantly, smFRET suggested that HIV-1 Env has spontaneous access to this intermediate conformational state and its occupancy increased in response to CD4 binding^22^. Our cryo-ET structures of asymmetric HIV-1 Env trimers bound to one and two CD4 molecules demonstrate that these intermediates are quite common and long-lived enough to be readily detected in our sampling of Env-CD4 complexes. The finding that these states are long-lived intermediates likely has consequences for antibody-mediated immune responses and vaccine immunogen design. Indeed, antibodies recognizing partially open conformational states, with the V1V2 loops of gp120 remaining at the apex of the trimer, have been identified^35,36^.

Our studies point to a stepwise model of HIV-1 binding to CD4^+^ cells early in the infection process (Figure 4o). Initial contact between HIV-1 virions and cells is mediated by binding of a single protomer to an elongated CD4 receptor molecule when the membranes are still ∼20 nm apart from each other. Engagement of a second and third CD4 molecule allows HIV-1 Env to move closer to the target membrane. This movement is facilitated by conformational changes within CD4 where the D3D4 domains align with the target membrane. However, even with these conformational changes the Env trimer bound to three CD4 molecules remains too far away to engage the membrane-embedded co-receptor (Fig. 4o). There are considerable steric constraints in the gp120-CD4 interaction that must be overcome for the Env trimer to bind the co-receptor. This could be achieved by either 1) shedding of the gp120-CD4 complex, or 2) membrane bending in the target cell. Future experiments with CCR5 embedded in target membranes will shed light on co-receptor binding and subsequent formation of the prehairpin intermediate. The experimental system described here will assist in characterizing these subsequent steps of the HIV-1 entry process.

## Acknowledgements

We thank Dr. Shenping Wu at the Yale CryoEM facility for her technical assistance. We thank Drs. James Binley, Heinrich Gottlinger, Richard Mulligan, Joseph Sodroski, and Zene Matsuda for plasmids. This work was supported by NIH grants R01 AI150560 to W.M. and R01AI143563 to M.B.Z. E.N. and M.W.G are supported by NIH T32AI055403. M.W.G. is a recipient of Gruber Science Fellowship. J.L. and J.W.B. Jr, are supported in part with federal funds from the National Cancer Institute, National Institutes of Health, under Contract No. 75N91019D00024/HHSN261201500003I.

## Author Contributions

W.L., E.N., and W.M. conceived experiments; W.L., E.N., J.W.B., J.D.L., J.R.G. M.B.Z, and P.D.U. produced HIV-1 and MLV viral particles; W.L. and M.W.G. collected cryo-ET data; W.L., M.W.G., E.N., and H.D.T. analyzed cryo-ET data; E.N. performed fusion kinetics experiments; Z.Q. performed the MDFF; E.N., W.L., and W.M. wrote original manuscript draft; all authors reviewed and edited manuscript; W.L. and W.M. supervised the work; W.M. acquired funding.

## Materials and Methods

### Plasmids, cell lines, and other reagents

HIV-1_BaL_/SUPT1-CCR5 cells (cell line ID #CLN204) ^37^ were cultured in RPMI-1640 media, supplemented with 10% fetal bovine serum (FBS), 100 U/mL penicillin/streptomycin, and 2 mM L-glutamine in the presence of 5% CO2. HEK293T and HIV-1_ADA.CM.755*_/HEK293T cells were cultured in DMEM media supplemented with 10% FBS, 100 U/mL penicillin/streptomycin, and 2 mM L-glutamine in the presence of 5% CO2. Cells were transfected at 60%–80% confluency and culture media was exchanged before transfection. HIV-1 GagPol was expressed by pCMV ΔR8.2 (Addgene plasmid # 12263). The protease mutations D25N and R57G were generated by overlapping PCR using pCMV ΔR8.2 as a template. The mutations were validated by Sanger sequencing. Human CD4 expressing vector pcDNA-hCD4 was kindly provided by Heinrich Gottlinger. The pCAGGS HIV-1_JR-FL_ gp160 expression plasmid was kindly provided by James Binley. The pCAGGS HIV-1_JR-FL_ Δ_CT_ gp160 was generated by introducing a stop codon to replace Gly/711 of gp160 in the pCAGGS HIV-1_JR-FL_ gp160 plasmid. The CypA-HiBit plasmid was derived from pEGFP-N1-CypA. To generate this plasmid, human CypA was cloned from HeLa cDNA and inserted into pEGFP-N1 using the EcoRI and BamHI sites. To generate CypA-HiBit, a synthetic oligo, which encoded the linker sequence “GSGSSGGGGSGGGGSSG” followed by the HiBit peptide “VSGWRLFKKIS,” was inserted to replace EGFP at the C-terminus of CypA using the BamHI and NotI sites. This construct is based on a similar design first utilized by Gregory Melikyan^38^. The pE7-NL4-3-Env plasmid was a kind gift from Joseph Sodroski. The pMX-puro PH-PLCΔLgBiT plasmid was kindly provided by Zene Matsuda^39^. The MLV GagPol plasmid was a kind gift from Richard Mulligan. The following reagents were obtained through the NIH HIV Reagent Program, Division of AIDS, NIAID, NIH: Indinavir sulfate, ARP-8145 and Ritonavir, ARP-4622, contributed by DAIDS/NIAID; Human CCR5 expression vector (pcCCR5), ARP-3325, and Human CXCR4 expression vector (pcCXCR4), ARP-3326, both contributed by Dr. Nathaniel Landau.

### Virus preparations for cryoET

HIV-1_BaL_ virus lot P4311 were produced by the HIV-1_BaL_/SUPT1-CCR5 cell line and prepared as previously described^19^. Briefly, up to 30 liters of cell culture were serially filtered using 5-µm capsule filters to remove the cells, followed by filtration using 0.45 µm capsule filters to remove cell debris and microvesicles. The filtrate was treated with a final concentration of 1LmM 2,2’-dithiodipyridine (Aldrithiol™-2, or AT-2). AT-2 eliminates retroviral infectivity by mediating covalent modification of free cysteines on internal viral proteins, including the zinc finger cysteines on the viral nucleocapsid protein. Even after AT-2 treatment, the envelope glycoproteins with disulfide-bonded cysteines on the virion surface retain the ability to bind to CD4, undergo conformational changes, and mediate membrane fusion^20,25^. HIV-1_BaL_ viruses were then purified by continuous-flow sucrose density-gradient ultracentrifugation.

Immature HIV-1_BaL_ viruses were produced by HIV-1_BaL_/SUPT1-CCR5 cell line treated with a final concentration of 1 µM Indinavir and 10 µM Ritonavir. Cells were treated for 2 days, after which the cell media was discarded and replaced with fresh media with the same concentrations of Indinavir and Ritonavir for two additional days. Cell culture media was then harvested and clarified by low speed spinning at 1500 rcf for 15 minutes, followed by filtration with 0.45 µm syringe filters (Pall Corporation). Filtered supernatants were loaded into 38.5 mL open-top ultracentrifuge tubes (Beckman Coulter) and underlayed with 5 mL of 15% sucrose in PBS. Viruses were then pelleted by ultracentrifugation at a maximum of 131,453 rcf using a Beckman SW28 swinging bucket rotor at 27000 rpm for 1 hour at 4°C. Supernatant was removed and viruses were resuspended in 1/1000 volume of PBS.

HIV-1_ADA.CM.755*_ was produced from ADA.CM.755*/293T cell line^30^ by transfection with 10 µg DNA per 10 cm plate of the HIV-1 GagPol plasmid pCMV ΔR8.2 using polycation polyethylenimine (PEI) (pH 7.0, 1 mg/mL). Cell culture media was collected two days after transfection, then clarified and filtered as above. Viruses were pelleted by ultracentrifugation and resuspended in PBS as above.

Immature HIV-1_ADA.CM.755*_viruses were produced from the ADA.CM.755*/293T cell line either by transfecting 10 µg DNA per 10 cm plate of pCMV ΔR8.2 in the presence of 1 µM Indinavir and 10 µM Ritonavir or by transfecting pCMV ΔR8.2 containing the protease mutations D25N or R57G. Viruses were collected after two days using the same methods as above.

MLV particles carrying CD4 receptor (MLV-CD4 VLPs) were produced by co-transfecting plasmids encoding MLV GagPol and CD4 at a 1:1 ratio into 293T cells using PEI. Viruses were collected after two days using the same methods as above.

### Cryo-electron tomography sample preparation

HIV-1_BaL_ particles and MLV-CD4 VLPs were mixed and incubated at room temperature for 30 minutes, after which a 6 nm gold tracer was added at a 1:3 ratio. 5 µL of the mixture was placed onto freshly glow discharged holey carbon grids (Quantifoil™ R 2/1 on 200 copper mesh) for 1 minute. Grids were blotted with filter paper and plunge frozen into liquid ethane using a homemade gravity-driven plunger apparatus. Frozen grids were stored in liquid nitrogen prior to imaging.

### Cryo-electron tomography data collection

Cryo-grids were imaged on a Titan Krios G4 Cryo-transmission electron microscope (cryo-TEM)(Thermo Fisher Scientific) operated at 300 kV using either a K2^®^ or K3^®^ direct electron detector (Gatan) in counting mode with a 20 eV energy slit and a Volta phase plate (VPP) ^40^. Tomographic tilt series between −51° and +51° were collected by using SerialEM^41^ using a 3° step size and a dose-symmetric scheme^42,43^. The nominal magnification for the K2^®^ direct electron detector was 105,000x, giving a pixel size of 1.333 Å, while for the K3^®^ direct electron detector it was 64,000x, giving a pixel size of 1.346 Å. The raw images were collected from single-axis tilt series with a cumulative dose of 123e/Å^2^. The defocus was set at −0.5 µm. Frames were motion-corrected using Motioncorr2^44^ to generate drift-corrected stack files, which were then aligned using gold fiducial makers using Etomo^43^. Weighted back projection and tomographic slices were visualized with IMOD.

### Cryo-electron tomography data analysis

Env-CD4 complexes at membrane-membrane interfaces were picked manually. Euler angles were determined based on the vector between two points, one on the HIV-1 particle membrane and the other on the MLV VLP membrane when the membranes were closest together.

Subtomograms were extracted for initial alignment. Subsequent processing was performed using I3^45^ with 4x and 2x binned tomograms. All the density maps were visualized and segmented in the UCSF ChimeraX^46^.

To quantify Env clustering, all Env trimers on virions were manually picked. The arc distance between any two Env trimers on each individual virion was calculated with a home-made script based on the approximate center of the virion and the Env positions after initial subtomogram averaging. The histogram of Env distances was plotted in GraphPad Prism 9.0.

### Fusion kinetics assay

A live-cell fusion assay using split nanoluciferase (nanoluc) complementation was developed to evaluate fusion kinetics of HIV-1 Env strains in real time. The split nanoluc system was used as previously described^39,47^. HIV-1 VLPs were prepared in HEK293E cells by co-transfecting HIV-1 GagPol, CyclophilinA-HiBiT, and the HIV-1 Env strain being tested or carrier DNA pcDNA 3.1 (Invitrogen) in a 1:1:1 ratio using 3 µL PEI per 1 µg DNA. Transfections were calculated using 12 µg of DNA per 10 cm plate. The CyclophilinA-HiBiT fusion protein is efficiently incorporated into viral particles as CyclophilinA binds to the HIV-1 capsid^48,49^. Cell culture media was collected approximately 48 hours after transfection and spun for 5 minutes at 551 rcf using a Sorvall 75006445 swinging bucket rotor at 1600 rpm to remove cell debris, followed by filtration with a 0.45 µm syringe filter (Pall Corporation). Filtered supernatants were loaded into open-top ultracentrifuge tubes (Beckman Coulter) and underlayed with 5 mL of 15% sucrose in PBS, then ultracentrifuged at a maximum of 131,453 rcf using a Beckman SW28 swinging bucket rotor at 27000 rpm for 1 hour at 4°C to pellet VLPs. The supernatants were discarded, and the pellets were resuspended in 1/100 volume of RPMI-1640 media supplemented with 10% FBS, 100 U/mL penicillin/streptomycin, and 2 mM L-glutamine. Resuspended pellets were incubated at room temperature for 30-60 minutes before being aliquoted and stored at −80°C. For preparation of immature HIV-1 VLPs, Indinavir and Ritonavir, at final concentrations of 1 µM and 10 µM respectively, were added to fresh media on HEK293E cells before transfection. Cells were co-transfected with HIV-1 GagPol, CyclophilinA-HiBiT, and the HIV-1 Env strain being tested as above. Supernatants were harvested after two days, after which cell debris was pelleted, supernatant was filtered, and VLPs were pelleted by ultracentrifugation through a 15% sucrose cushion as with the mature HIV-1 VLPs above. Supernatants were removed after ultracentrifugation and immature VLPs were resuspended in RPMI-1640 medium supplemented with 10% FBS, 100 U/mL penicillin/streptomycin, 2 mM L-glutamine, and also with 1 µM Indinavir and 10 µM Ritonavir. Resuspended pellets were incubated at room temperature for 30-60 minutes before being aliquoted and stored at −80°C as above. All VLP volumes were calculated and normalized to 1.4 x 10^7^ RLU using the Nano-Glo^®^ HiBiT Lytic Detection System kit (Promega).

To prepare target cells for the virus-cell fusion assay, HEK293E cells were transfected with PH-LgBiT, CD4, and CCR5 or CXCR4 plasmids at a 1:1:1 ratio using PEI at a 3 µL per 1 µg DNA ratio and using 12 µg total DNA per 10 cm plate. The PH domain from phospholipase C-delta (PLC-δ) ^50^ localizes LgBiT to the plasma membrane of the target cells, thereby allowing for immediate reconstitution of HiBiT and LgBiT into the complete nanoluc protein upon VLP fusion. At 24 hours after target cell transfection, cells were resuspended in their same transfection media. A master mix of transfected cells with 1x Endurazine™ (Promega) and 1 µM DrkBiT peptide (VSGWALFKKIS, synthesized by GenScript at >95% purity) was then prepared and added to wells in triplicate on four white 96-well plates (Greiner Bio-One). The membrane-permeable nanoluc substrate Endurazine™ (Promega) was included in the media on the target cells to allow for immediate light production upon reconstitution of the nanoluc activity. The membrane-impermeable DrkBiT peptide is a competitive binder of LgBiT. DrkBiT binding does not reconstitute the nanoluc protein, thus reducing background from free LgBiT in the media^39^. Plates, VLPs, pipette, and tips were then pre-incubated in a cold room at 4°C for 10-30 minutes. Normalized VLP volumes were added to the transfected cell master mix on the cold plates in the cold room at 4°C. All four white 96-well plates were exact replicates for thermal control during kinetics measurements. Plates were kept on ice as they were transferred to a pre-cooled centrifuge for a 2-hour spinoculation at 1200 rcf and 12°C. After the 2-hour spinoculation, plate numbers 2, 3, and 4 were placed in a 37°C incubator and time was noted as 0 minutes.

Meanwhile, plate 1 was read by the luminometer as the 0-minute time point. Plate 1 was then moved to the 37°C incubator. Plate 2 was measured on the luminometer at 10 minutes, then returned to the incubator. This process was repeated for plates 3 and 4 for the 20-minute and 40-minute time points. Plate 1 was read again for the 60-minute time point. Fusion kinetics data were analyzed by dividing each sample reading by the average of the master mix only negative triplicate readings on each plate. Each plate reading was analyzed independently. The data were then plotted in GraphPad Prism 9.0.

### Molecular dynamics flexible fitting

The cryo-EM density map obtained from subtomogram averaging was low-pass filtered to 15 Å resolution. The fully open atomic model of CD4-bound B41 SOSIP (PDB: 5VN3) and the partially open atomic model of CD4-bound BG505 SOSIP (PDB: 6CM3) were first fit into the density map by rigid-body fitting in Chimera^51^. To build the initial atomic models of Env-CD4 complex, the full-length CD4 molecules from CD4-gp120-CCR5 complex (PDB: 6MET) were added to both SOSIP trimers 5VN3 and 6CM3 by aligning and superimposing the gp120 subunits. The coordinates of the initial atomic models were saved relative to the density map.

The initial models were then prepared using the Visual Molecular Dynamics program (VMD) ^52^ to generate the constrains to preserve the secondary structure, chirality, and cis-peptide bonds. The Molecular Dynamics Flexible Fitting (MDFF) of Env-CD4 complexes were performed in NAMD 3.0 Alpha^53^ with CHARMM36^54^ force-field parameters, *in vacuo* with isothermal conditions at 300 K. The fitting of the combined partially open model took 1.5 nanoseconds with the MDFF force scaling factor ξ = 0.1 followed by an energy minimization step of 0.05 picoseconds with the scaling factor ξ = 0.5. The combined fully open model used 2.5 nanoseconds with ξ = 0.1 followed by an energy minimization step of 0.05 picoseconds of fitting with the scaling factor ξ = 0.5.

## Figure legends

**Extended Data Fig. 1.**
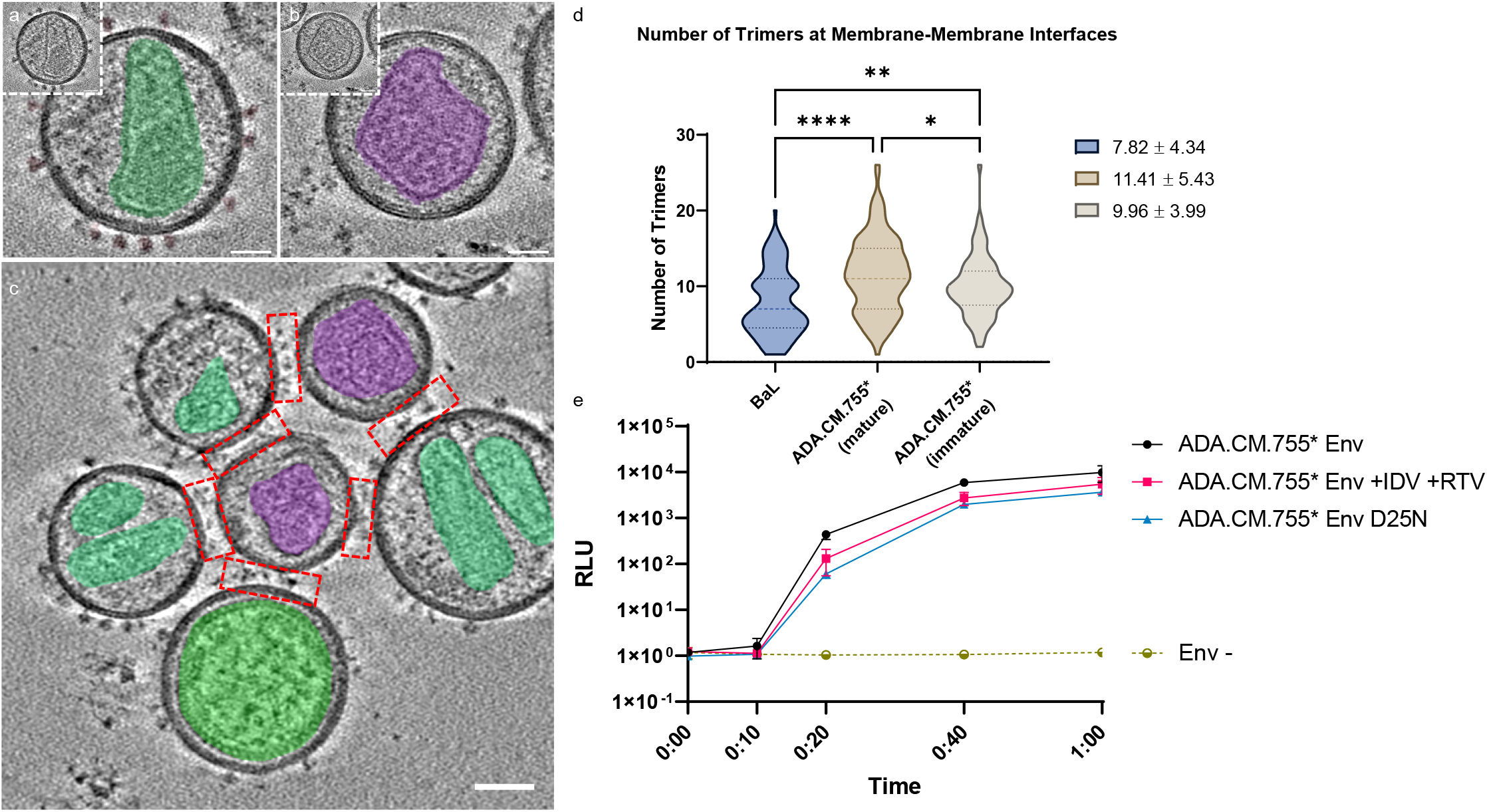
HIV-1 Env clustering and fusion kinetics are unaffected in HIV-1 particles featuring a high number of Env trimers lacking the C-tail. **a** Representative images of the HIV-1 viral particle produced from the cell line expressing Env_ADA.CM.755*_. The capsid is highlighted in green. Scale bar = 25 nm. **b** Representative images of the MLV VLP carrying CD4 and CCR5. The capsid is highlighted in purple. Scale bar = 25 nm. **c** A representative image of membrane-membrane interfaces in cryo-tomograms. HIV and MLV capsids are highlighted in green and purple, respectively. Scale bar = 50 nm. **d** Violin plot of the number of Env trimers present at membrane-membrane interfaces for HIV-1_BaL_ (blue), mature HIV-1 Env_ADA.CM.755*_ (brown), and immature HIV-1_ADA.CM.755*_ (beige) with MLV-CD4-CCR5 VLPs. P-values * =0.0315, ** =0.0021, **** = <0.0001. **e** Fusion kinetics of mature HIV-1_ADA.CM.755*_ (black), and of particles made immature in presence of protease inhibitors IDV and RTV (red) or with the PR-mutation (blue). Two independent experiments were performed in triplicate and data sets for each sample were combined and plotted. Error bars are SD.

**Extended Data Fig. 2.**
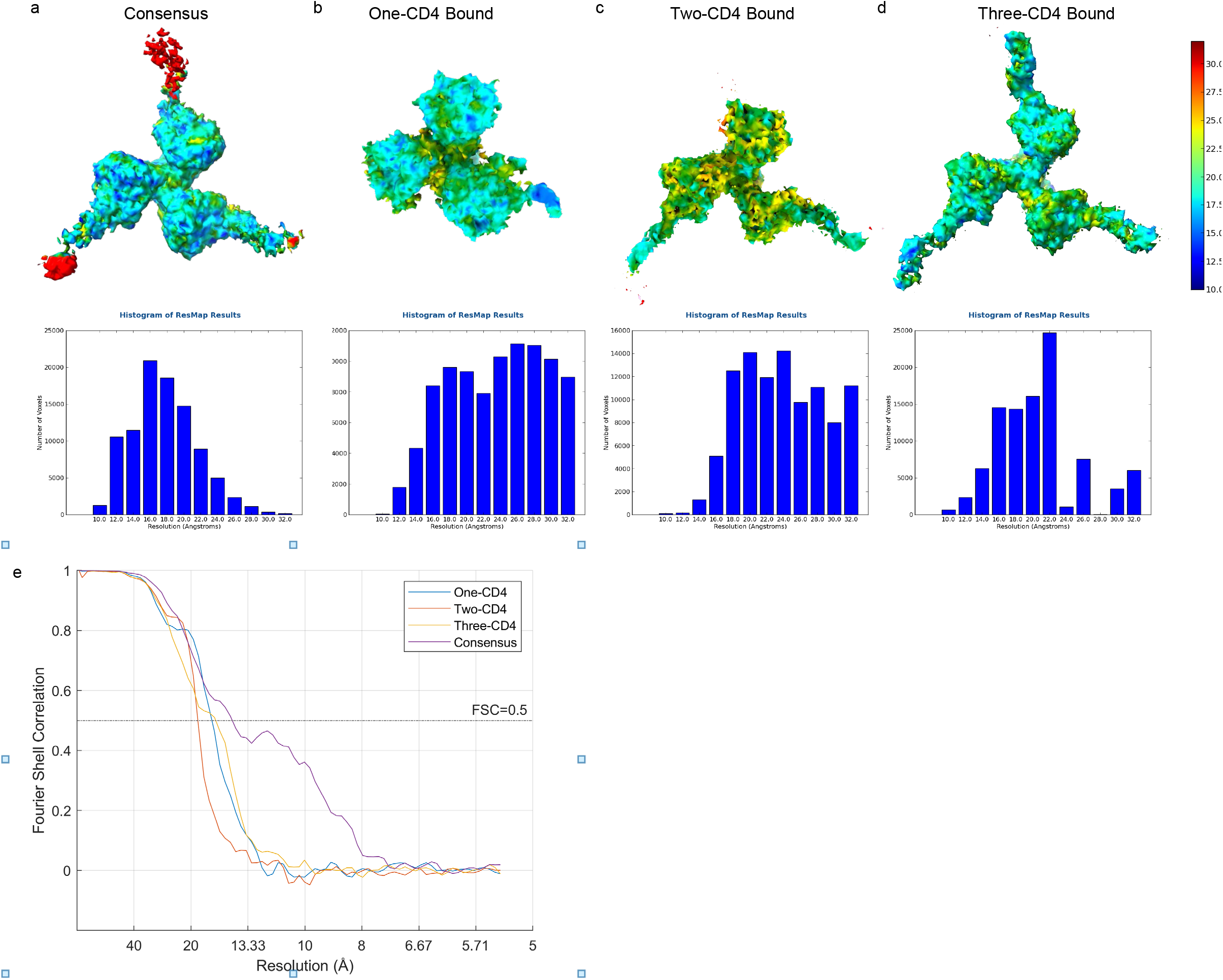
Resolution assessment of subtomogram averaged structures. **a-d** Local resolution estimation using ResMap software for the structures of the Env-CD4 complex. The structures and ResMap histograms of the consensus average (a), Env trimer bound to one CD4 molecule (b), Env trimer bound to two CD4 molecules (c), and Env trimer bound to three CD4 molecules (d) are shown. Subtomogram averages are colored according to the local resolution as indicated on the scale on the right. **e** Fourier shell correlation (FSC) curves for cryo-ET masked subtomogram averages with FSC = 0.5 as the cutoff value.

**Extended Data Fig. 3.**
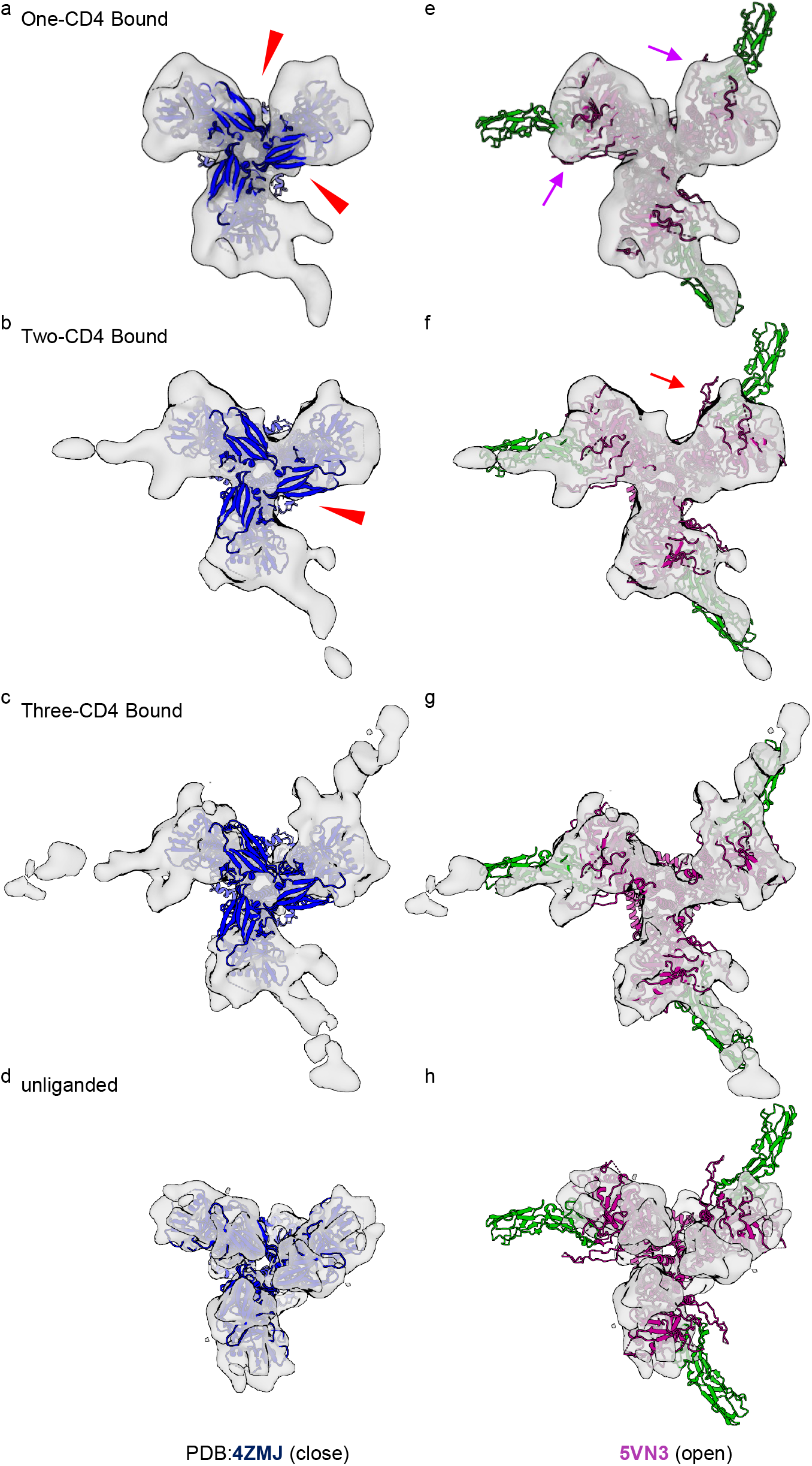
HIV-1 Env trimers bound to one and two CD4 molecules are asymmetric with protomers adopting distinct conformational states. **a-h** Rigid fitting of atomic structures of Env in a closed conformation (PDB:4ZMJ^55^)(a-d) and an open conformation (PDB:5VN3)(e-h) into the density maps of Env bound to one CD4 (a, e), two CD4 (b, f), three CD4 molecules (c, g) and unliganded Env (EMDB:21412) (d, h). Missing densities at the apex region of non-CD4 bound protomers are indicated by red arrowheads. Weak densities of the V1V2 loops on the non-CD4 bound protomers in the Env trimer bound to one CD4 molecule are indicated by purple arrows (e). Lack of density for the outward projecting V1V2 loops on the non-CD4 bound protomer in the Env trimer bound to two CD4 molecules is indicated by a red arrow (f).

**Extended Data Fig. 4.**
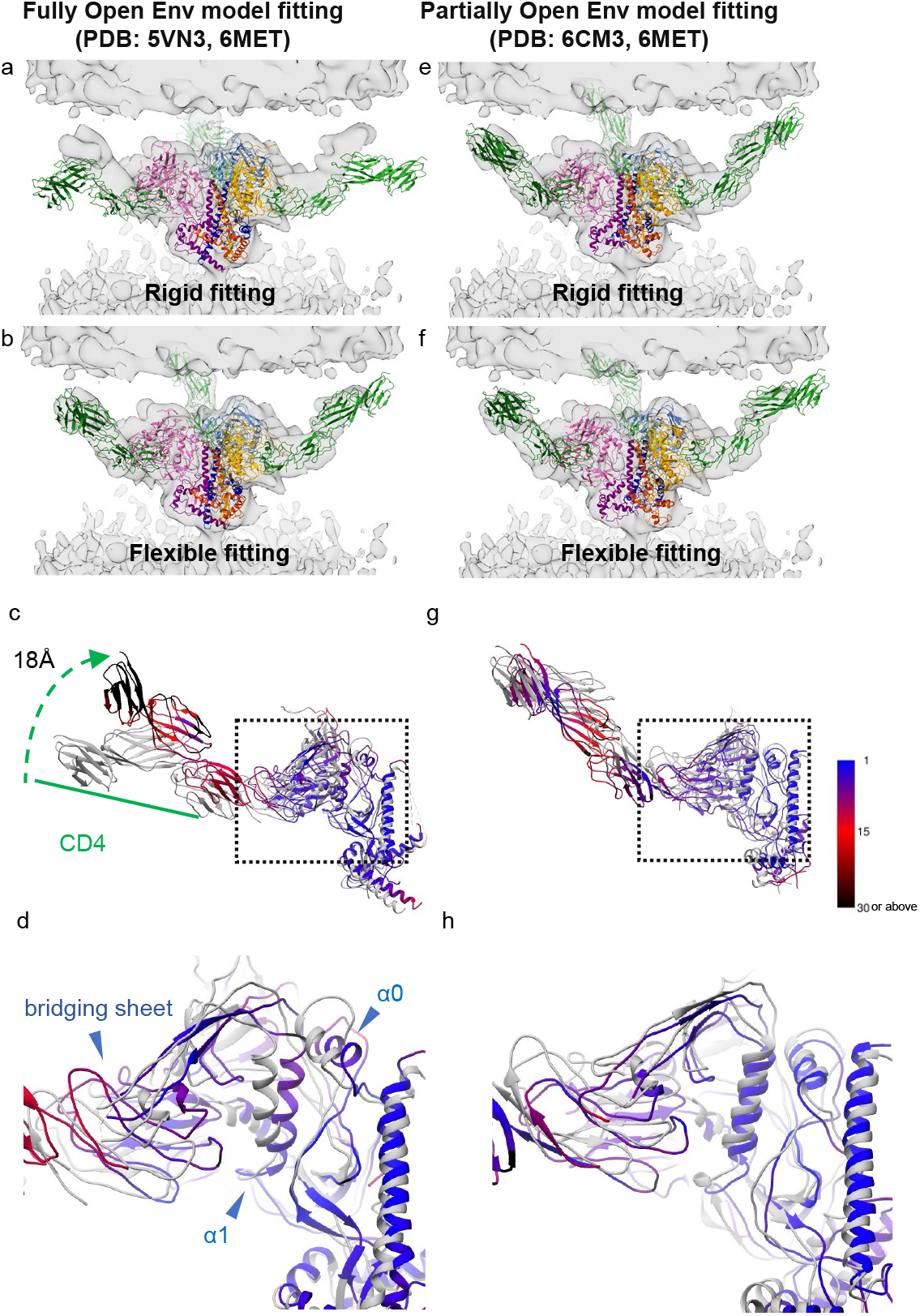
Molecular dynamics flexible fitting suggests that the Env trimer bound to three CD4 molecules is in a partially open conformation. **a** Rigid body fitting of the combined atomic model of full-length CD4 molecules (from PDB: 6MET) superimposed on fully open Env (from PDB: 5VN3) into the density map of Env bound to three CD4 molecules. **b** Molecular dynamics flexible fitting (MDFF) of the combined atomic model in panel (a) into the same density map of Env bound to three CD4 molecules. **c, d** Atomic models showing the positions of CD4 (c) and gp120 (d) before (grey) and after (colored) MDFF in panel (b), respectively. Color gradient indicates the amount of movement between the two structures as indicated on the right. **e** Rigid body fitting of the combined atomic model of full-length CD4 molecules (from PDB: 6MET) superimposed on partially open Env (from PDB:6CM3) into the same density map of Env bound to three CD4 molecules. **f** MDFF of the combined atomic model in panel (e) into the same density map of Env bound to three CD4 molecules. **g, h** Atomic models showing the positions of CD4 (g) and gp120 (h) before (grey) and after (colored) the MDFF in panel (f), respectively. Color gradient as in (c) and (d).

**Extended Data Table 1.** Shifts in the center of mass for the domains of each Env protomer and CD4 molecule after flexible fitting (Å). Table of the shifts in the center of mass for the domains of Env protomers and CD4 after the MDFF analysis in Extended Data Fig. 4.

| Domains<br>5VN3<br>6CM3 | Protomer1 | Protomer2 | Protomer3 |
| --- | --- | --- | --- |
| gp120 | 2.36 | 3.11 | 2.39 |
|  | 1.16 | 2.89 | 1.77 |
| gp41 | 1.57 | 2.37 | 3.08 |
|  | 2.13 | 3.36 | 1.84 |
| CD4 | 18.74 | 16.56 | 16.49 |
|  | 7.46 | 8.31 | 6.75 |
| gp120 α0 Helix | 3.08 | 4.89 | 3.70 |
|  | 2.10 | 1.39 | 2.77 |
| gp120 α1 Helix | 3.82 | 4.75 | 3.51 |
|  | 1.63 | 2.97 | 1.95 |
| bridging sheet | 5.53 | 7.06 | 7.60 |
|  | 1.47 | 4.59 | 3.96 |
| CD4 D1 | 7.12 | 5.44 | 5.90 |
|  | 4.64 | 5.71 | 3.92 |
| CD4 D2 | 14.93 | 12.09 | 11.72 |
|  | 9.29 | 11.52 | 9.00 |
| CD4 D3 | 22.72 | 19.94 | 20.57 |
|  | 10.86 | 12.92 | 10.58 |
| CD4 D4 | 34.86 | 36.09 | 32.42 |
|  | 9.00 | 8.39 | 8.77 |

**Extended Video 1. Molecular dynamics flexible fitting suggests that the Env trimer bound to three CD4 molecules is in a partially open conformation**. Molecular dynamics flexible fitting (MDFF) of the combined atomic model of full-length CD4 molecules (from PDB: 6MET) superimposed on fully open Env (from PDB: 5VN3) into the density map of Env bound to three CD4 molecules.

**Extended Video 2. Molecular dynamics flexible fitting suggests that the Env trimer bound to three CD4 molecules is in a partially open conformation**. Molecular dynamics flexible fitting (MDFF) of the combined atomic model of full-length CD4 molecules (from PDB: 6MET) superimposed on partially open Env (from PDB: 6CM3) into the density map of Env bound to three CD4 molecules.

